# Dynamic reorganization of nuclear architecture during human cardiogenesis

**DOI:** 10.1101/222877

**Authors:** Paul A. Fields, Vijay Ramani, Giancarlo Bonora, Gurkan Yardimci, Alessandro Bertero, Hans Reinecke, Lil Pabon, William S. Noble, Jay Shendure, Charles E. Murry

## Abstract

While chromosomal architecture varies among cell types, little is known about how this organization is established or its role in development. We integrated Hi-C, RNA-seq and ATAC-seq during cardiac differentiation from human pluripotent stem cells to generate a comprehensive profile of chromosomal architecture. We identified active and repressive domains that are dynamic during cardiogenesis and recapitulate *in vivo* cardiomyocytes. During differentiation, heterochromatic regions condense in *cis*. In contrast, many cardiac-specific genes, such as *TTN* (titin), decompact and transition to an active compartment coincident with upregulation. Moreover, we identify a network of genes, including *TTN*, that share the heart-specific splicing factor, RBM20, and become associated in *trans* during differentiation, suggesting the existence of a 3D nuclear splicing factory. Our results demonstrate both the dynamic nature in nuclear architecture and provide insights into how developmental genes are coordinately regulated.

**One Sentence Summary:** The three-dimensional structure of the human genome is dynamically regulated both globally and locally during cardiogenesis.

## Main Text

Human development requires precise temporal and spatial regulation of gene activity. Congenital heart disease, the most common birth defect, is often caused by mutations in transcription factors leading to abnormal gene expression (*1, 2*). Even today, the mechanism of how these signaling pathways control lineage specification is largely unknown. Chromosome conformation capture technologies such as Hi-C have shown that the three-dimensional organization of the genome is dynamic during development, and abnormal chromosomal architecture has been implicated in a range of diseases (*3–6*). We have previously shown that epigenetic modifications are dynamically regulated during cardiomyocyte differentiation (*7*), and that integration of epigenetic data defines general patterns of gene regulation during differentiation. To gain insights into the role of nuclear architecture in cardiac development we combined *in situ* DNase Hi-C (*8*), RNA-seq and ATAC-seq during cardiac differentiation from human pluripotent stem cells (hPSCs) (*9*). We show that cardiac development genes are regulated at the level of nuclear architecture and that these structural dynamics influence gene regulation transcriptionally. Because the nuclear architecture of hPSC-cardiomyocytes closely resembles that of human fetal cardiomyocytes, these dynamics likely reflect early steps in human cardiogenesis.

We generated highly pure populations of cardiomyocytes (CMs) from karyotypically normal undifferentiated RUES2 hESCs (Fig. 1A, S1A-C, >90% cTnT+). During differentiation, these cells pass through stages representative of early development including mesoderm (MES), and cardiac progenitor (CP), before reaching definitive CMs (*7, 9*) (fig. S1D). We performed *in situ* DNase Hi-C (*8*) on these stages of differentiation with two independent biological replicates, along with two fetal heart samples (table S1). Chromosome-wide contact maps demonstrate the expected checkerboard pattern, indicative of local compartmentalization (topology-associated domains, or TADs) and longer range compartmentalization (A/B compartments [see below]; Fig. 1B). Genome-wide contact maps between whole chromosomes demonstrate that smaller and larger chromosomes tend to self-associate. Few whole chromosome changes are observed across time points save for a shift in chromosome 15, from associating with larger chromosomes in earlier time points, to smaller chromosomes in CMs, matching data from the fetal hearts (fig. S2).

**Fig. 1:**
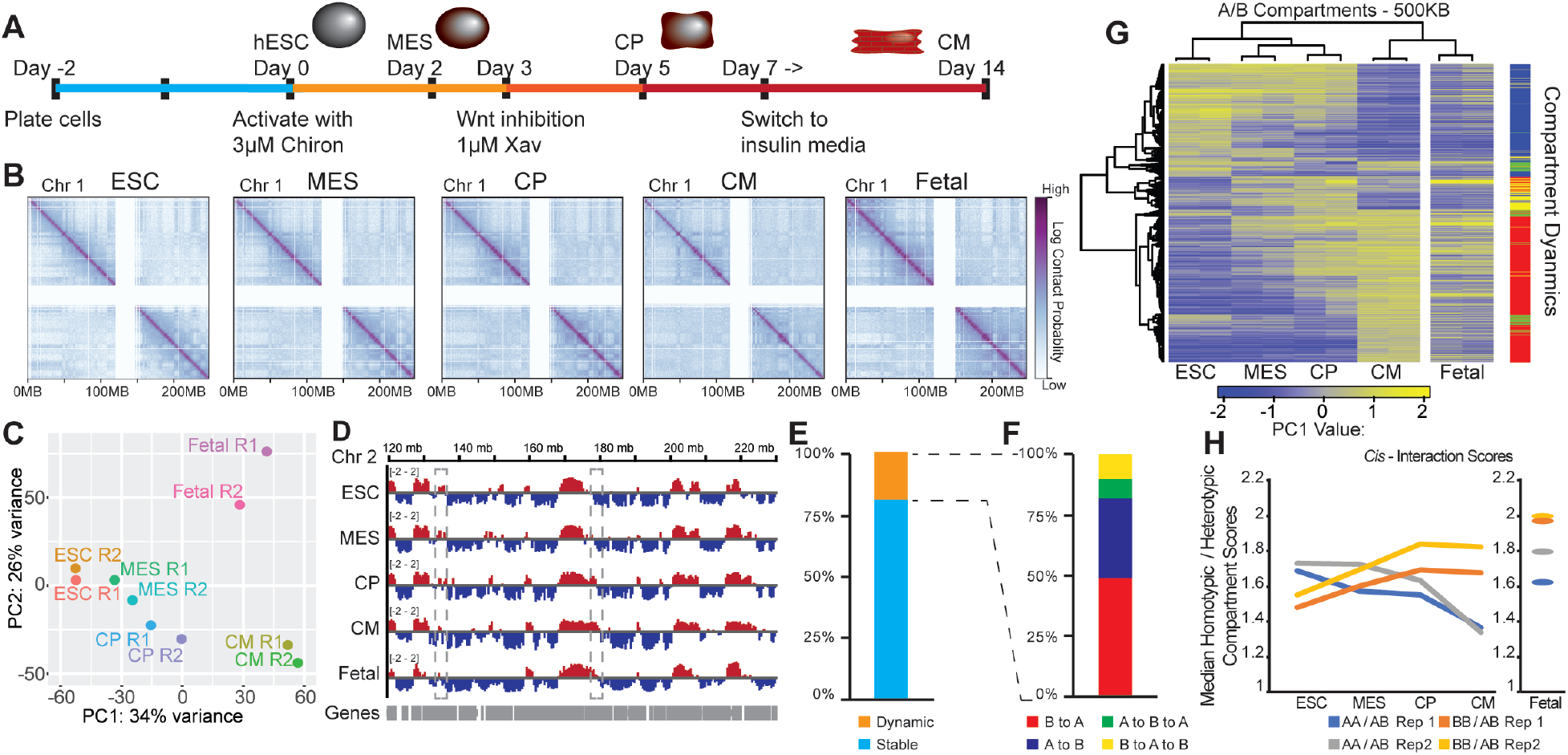
(A) Schematic of the cardiomyocyte differentiation. (B) Log transformed contact maps of chr 1. (C) PCA plot of PC1 scores on the contact matrices. (D) PC1 scores for a region of chr 2, grey boxes highlight regions A to B and B to A. (E) Genomic regions divided by stable and dynamic A/B compartment (ANOVA p<0.05). (F) Dynamic regions divided by types of compartment transitions. (G) Heatmap of the PC1 scores of the dynamic regions. Clustering of rows based on the four time points of differentiation. Dendrogram of columns was ordered to match the temporal status of differentiation. (H) Median value of A-A and B-B interactions divided by A-B across differentiation and in fetal heart for *cis* contacts.

Using eigenvalue decomposition of our contact maps, we separated each chromosome into A/B compartments, which reflect regions of active and repressive chromatin, respectively (*10*). Principle component analysis (PCA) of the A/B scores at 500kb resolution separates the samples by time point over PC1, while PC2 likely separates samples by cardiomyocyte purity (Fig. 1C). Early fetal hearts, while consisting of ~70% cardiomyocytes (*11, 12*), include other cell types such as fibroblasts. Overall the genome is split into ~50% A, ~50% B compartments at each time point (fig. S3A), and the majority of these compartmental assignments are invariant during differentiation (Figs. 1D-E). However, we find that 19% of the genome change compartment during differentiation, and that clustering of dynamic regions recapitulates the differentiation trajectory (Fig. 1G). Most of these changes are unidirectional, either from B to A or A to B. A small subset (approximately 4% of the genome) exhibits a transitory switch, either A-B-A or B-A-B (Fig. 1F). Less than 1% show three switches (B-A-B-A or A-B-A-B) and are combined with the B-A and A-B subsets in subsequent analysis. Together these data show that A/B compartments are dynamic during cardiac differentiation, and that these changes are validated by analyses of fetal hearts.

To examine the dynamic regions, we generated differential contact maps between CMs and hESCs (fig. S3B). We observe that many of the strongest gains in long range interactions are associated with the B compartment. Genome-wide analysis shows that stronger intra-chromosomal (*cis*) contacts occur between homotypic regions (A-A or B-B compartments), compared to between heterotypic regions (A-B) (fig. S3B). In hESCs, we observe that average *cis* interaction signal between A compartments is strongest, while during differentiation this switches to favor signal between B compartments—a trend supported by patterns in fetal heart (Figs.1H, S3C). In contrast, inter-chromosomal interactions (*trans*) are consistent over differentiation and favor A-A interactions (figs. S3D,E). Together these observations suggest a model whereby during differentiation heterochromatic regions pack more tightly, specifically within chromosome territories in *cis*, while inter-chromosomal interactions are most likely to occur between active regions.

To investigate those genes that may be drivers of compartment changes, we performed RNA-seq on the same stages of differentiation, along with the fetal hearts (table S2). Similar to Hi-C, PCA separates RNA samples by differentiation state and cardiomyocyte purity (Fig. 2A). We find that ~7500 genes are differentially regulated during differentiation (Figs. 2B, S4A). In contrast to Hi-C most of the dynamic changes are cell-type specific. While RNA-seq reveals similarities between hPSC-CMs and the fetal heart (Fig. 2B), the fetal heart has an increase in the ratio of adult to fetal myosin heavy chain genes (*MYH7* and *MYH6*), suggesting that these mid-gestational cardiomyocytes are more developmentally advanced. Additionally, fetal hearts contain cardiac fibroblast transcripts such as *POSTN* and endothelial transcripts such as *PECAM1* and *CDH5*, as would be expected from a multicellular tissue (*13, 14*) (fig. S4A). Genes that are upregulated in CPs and CMs are enriched in regions that transition B to A, and underrepresented in A to B transitioning regions (Fig. 2C). Gene ontology (GO) analysis reflects that genes upregulated in CMs are enriched in categories related to metabolism along with development and cardiac function (table S3). Focusing this analysis on those genes in regions that transition from B to A highly enriches for genes involved in heart development, such as the structural gene alpha-actinin (*ACTN2*) (Fig. 2D,E, table S4). This suggests that many heart development genes may initially be sequestered in the B compartment and move to A upon activation. Strikingly, even though only 4% of the genome shows 2-step dynamics (A-B-A or B-A-B; Fig. 1G), genes that are transiently in the A compartment show peak expression in the CP stage (Fig. 2C). Among these, *BMPER* and *CXCR4* are the highest expressed in CPs, both of which have important roles in cardiomyocyte function (*15, 16*) (fig. S4B). Gene repression is also associated with dynamic compartments. For instance, the mesoderm regulator *EOMES* is initially in A compartment (but surrounded by B region) and switches to B upon downregulation at the CP stage (fig. S4C). Taken together, these observations show that genomic regions dynamically transitioning between compartments occur coincident with transcriptional regulation and the observed patterns are consistent between stem cell derivatives and the fetal heart, suggesting important developmental control.

**Fig 2:**
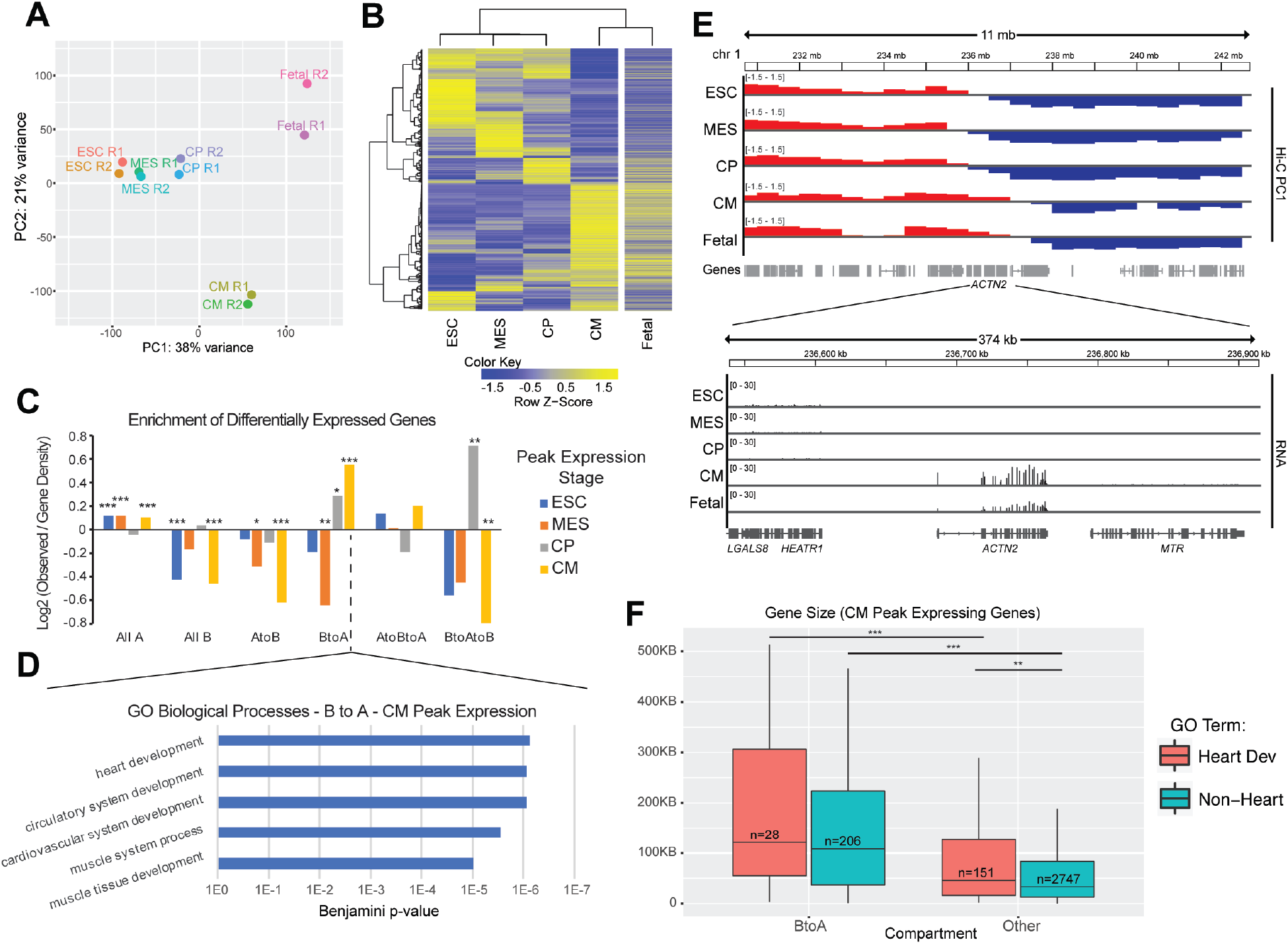
(A) PCA plot on the replicates of the RNA-seq samples. (B) Heatmap of the differentially expressed genes during differentiation, clustering of rows based on differentiation time points. (C) Enrichment of differentially regulated genes by time point of peak expression against A/B compartment dynamics. Log2 values are observed/gene density (p-value chi-sq test, *<0.05, **<0.01, ***<0.001). (D) GO term enrichment in CM peak expression genes in B to A compartments. (E) Gene track of PC1 and RNA-seq reads of *ACTN2*. (F) Boxplot of gene size of upregulated genes peaking in CM stage subdividing either B to A compartment or heart development genes (GO term).

Given the strong enrichment of CM-upregulated genes in the regions of the genome that transition from B to A, we sought to determine whether there are specific characteristics of these genes relative to other upregulated genes. Regions that transition from B to A have lower gene density (fig. S4D), suggesting these genes may be more isolated and exert a stronger influence on their region of the genome. We find that upregulated genes in B to A regions are larger than other upregulated genes (Fig. 2F). While upregulated genes are not more isolated (start site to start site) when normalized for gene size, heart development genes exhibit greater promoter isolation relative to other genes (fig. S4E,F), suggesting that isolation of cardiac development genes may be a mechanism to allow for individual regulation. Using expression data from 37 adult tissues (*17*) we find that those genes that are upregulated and go from B to A are more heart-enriched as determined by a higher expression rank in heart tissue compared to other upregulated genes (Fig. S4G). This supports a model by which large, lineage specific, cardiac genes are initially insulated in B compartment and transition to A upon activation. Further, the promoter isolation of developmental genes may allow for individual regulation.

While A/B compartments reflect the global organization of the genome across full chromosomes, local organization can be summarized as topologically associated domains (TADs). We utilized the directionality index (*18*) to call TADs. Given that TAD boundaries cluster by replicate (fig. S5A), we used TAD-calls on the merged samples for each condition for subsequent analyses. Most TAD boundaries are present in all four time points and therefore invariant over differentiation (fig. S5B). TADs in the A compartment are greater in number and smaller in size (fig. S5C), which is likely necessary for proper gene regulation in gene-dense areas. Comparing hESCs to CMs, hESC-specific boundaries are enriched within regions that are always in B or transit from A to B, consistent with our previous analyses suggesting that B-B domains compact during differentiation (fig. S5D). Boundaries that are gained during differentiation are associated with activation of the nearest gene (fig. S5E). As one example, the locus of *CAMK2D*, a kinase that regulates cardiac contractility and hypertrophy, changes local TAD structure coincident with upregulation of transcription (fig. S5F). While the majority of TAD boundaries are invariant, a substantial fraction change during differentiation and occur independent of A/B compartment changes. Neo-TAD boundaries occur adjacent to upregulated genes but separate from B to A dynamics, while loss of TAD boundaries occur preferentially within B compartment and supports a model where TAD boundaries are lost during heterochromatin compaction.

To determine whether changes in higher order organization occur coincident with local changes, we performed ATAC-seq to measure local accessibility across differentiation (table S5). Similar to our previous study using DNase hypersensitivity peaks (*19*), we find a greater number of peaks in hESC and MES compared to CM (Fig. 3A). Clustering pairs replicates, and orders the samples by differentiation state (fig. S6A). During differentiation, there is an increase in the fraction of peaks in A compartment (Fig. 3B, S6B,C), consistent with the Hi-C data, and supporting the model that heterochromatic regions becomes more condensed. Stage-specific peaks are on average more distal to TSSs than less dynamic peaks, consistent with enhancer activity (fig. S6D). Similar to the gene expression data, we find that CM-specific ATAC-seq peaks are enriched in regions that transition from B to A and are depleted in A to B regions. hESC-specific peaks are enriched in regions that are constitutively B compartment, consistent with the overall decrease in B region accessibility (Fig. 3C). Stage-specific peaks are enriched in motifs corresponding to developmental transcription factors (fig. S6E), in agreement with another recent study (*20*). Interrogating the regions that transition from B to A, we find that these regions are enriched for peaks present in CP and CM stages and significantly depleted in constitutive peaks (Fig. 3D). The top motifs within these peaks matched GATA and NKX motifs, both important cardiogenic transcription factors (Fig. 3E). GATA is a pioneer transcription factor and thus may play a pivotal role in opening up chromatin and driving the transition from B to A compartment(*21*). One such example is the gene *NEBL*, which encodes the sarcomeric protein nebulette, and is often mutated in familial dilated cardiomyopathies (*22*). The *NEBL* locus transitions from B to A compartment in CM stage, following increased accessiblity of a GATA motif in its promoter and coincident with large-scale transcriptional activation in the CM stage (Fig. 3F). Thus, local changes in chromatin accessibility occur coincident with changes in RNA expression and large scale genome organization.

**Fig 3:**
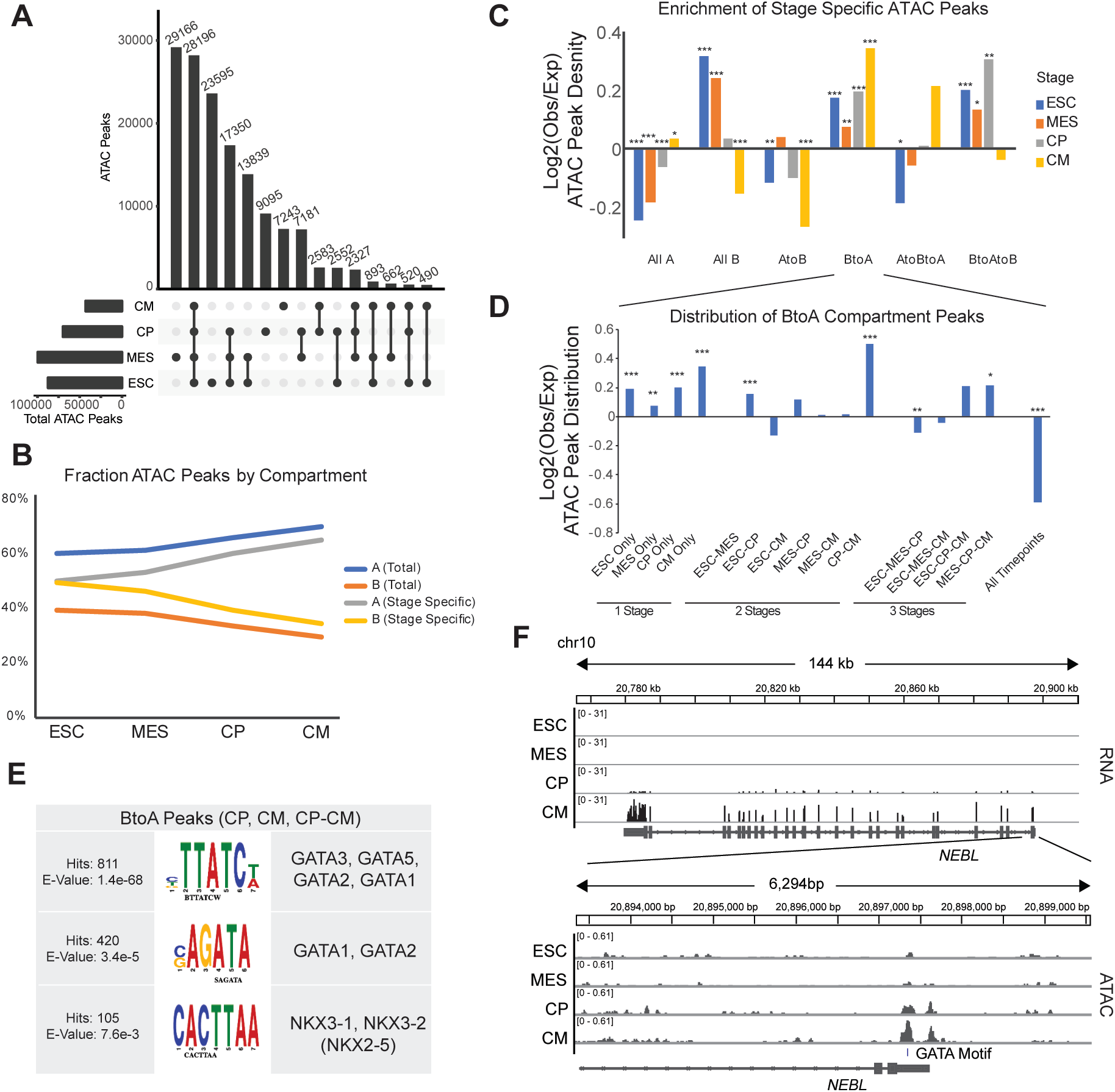
(A) ATAC peaks across differentiation. (B) Fraction of ATAC peaks divided by A and B compartment. (C) Enrichment of stage specific ATAC peaks within A/B compartments. Log2 Observed/ATAC peak density. (D) Enrichment of peaks within the B to A compartment bins by stage specificity, normalized to ATAC peak density (p-value chi-sq test, *<0.05, **<0.01, ***<0.001). (E) Motifs from DREME and TOMTOM within CP and CM specific peaks in B to A compartment bins. (F) Gene track of ATAC and RNA-seq reads for *NEBL* gene.

A major finding from our analyses is that regulation of large, developmental genes specifically relies on structural changes in genome organization. A prominent example is titin, an essential cardiac protein, which when mutated is the leading cause of dilated cardiomyopathy (*23*). The *TTN* locus transitions from B to A, coincident with an opening up of the local chromatin structure and upregulation of transcription (Figs. 4A, S7A). There is a corresponding increase in accessibility at the full length promoter of *TTN* as well as an internal promoter (*24*) (Figure 4B). Interestingly, we also observe three peaks of accessibility within *TTN* that decrease during differentiation (one 1KB downstream, another ~100KB into the gene and the third at the TTS), all of which overlap CTCF peaks (*25*). This is suggestive of a physically proximal chromatin hub at this locus in hESCs (*26*). Indeed, focusing on *cis* interactions upstream and downstream of *TTN’s* TSS, we identified a decrease in downstream signal during differentiation, with little change upstream (fig. S7B). Together, these data support a model where in hESCs the *TTN* locus is tightly compact, and during differentiation the locus opens up, transitions from B to A compartments, and activates transcription.

**Fig 4:**
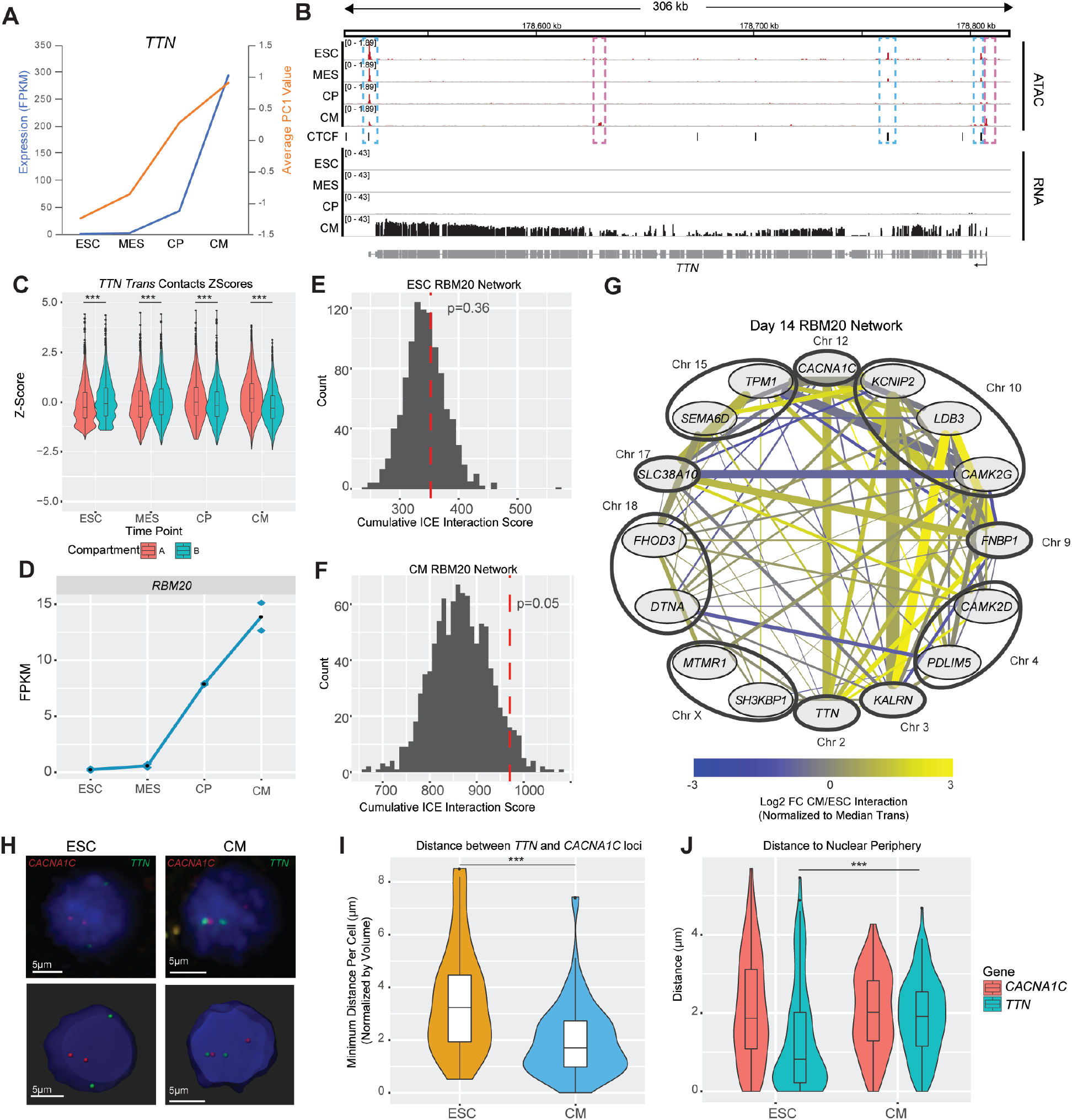
(A) PC1 value of the *TTN* locus and the FPKM expression levels across differentiation. (B) Gene track of the *TTN* gene with both ATAC and RNA signal and CTCF peaks. (C) Boxplot of the *trans* contacts of *TTN* by compartment, z-scored by time point at 500KB resolution. (D) Expression plot of *RBM20* across differentiation. (E-F) Cumulative score of association between upregulated RBM20 target genes in ESC (E) and CM (F) indicated by red line, background model is 1000 random permutation of selected genes from the same chromosomes. (G) Network of upregulated RBM20 genes, line thickness proportional to contact score, only scores greater than median *Trans* contact displayed. (H) Representative 3D FISH images in ESCs and CMs, and after spot calling processing. (I) Boxplot of normalized minimum distance per cell between a *TTN* locus and *CACN1AC* locus. (J) Boxplot of the distance from each locus to the nuclear periphery. All p-values, Wilcoxon test, ***<0.001.

Interactions between genomic regions in *trans* might involve important gene regulatory complexes such as transcriptional or splicing machinery. *TTN’s* move to the A compartment during cardiogenesis prompted us to explore its *trans* interactions in detail. During cardiac differentiation there is a switch in *TTN’s trans* interactions from an enrichment with B compartment to A (Fig. 4C). The regions within the enriched *trans* contacts in the CM stage include 279 genes that are also upregulated during the differentiation, including genes involved in cell surface signaling and muscle contraction (table S6). Two of those TTN-interacting cardiac genes are *CACNA1C* (which encodes a component of the L-type calcium channel; Chr 12) and *KCNIP2* (which encodes a voltage-gated potassium channel; Chr 10) (fig. S7C). All three of these genes are subject to alternative splicing, a process regulated by the cardiac-specific protein RBM20 (*27*). RBM20 promotes exon exclusion, and mutations in RBM20 lead to cardiomyopathies (*28*). We focused on a set of 16 genes, that show both conserved splicing regulation by RBM20 (*27*) and are upregulated during cardiac differentiation. These 16 genes show a significant Hi-C association in CMs but not in ESCs, with most pairwise interactions showing increased contacts (Figs. 4E-G). To validate the interaction between the *TTN* and *CACNA1C* loci, we performed 3D DNA FISH (Fig. 4H). We find that the minimum normalized distance between a *TTN* locus and *CACNA1C* locus is significantly less in CMs (Fig. 4I). Moreover, during differentiation *TTN* moves significantly further away from the nuclear periphery, suggesting that this gene might be found in a heterochromatic lamin-associated domain in hESCs (*29*), and in agreement with its B to A transition during cardiogenesis. Conversely, the constitutively A compartment gene *CACNA1C* is found at the same distance from the nuclear periphery in hESCs and CM (Fig. 4J). Because previous studies have shown that the *TTN* locus and RBM20 protein co-localize in two nuclear foci in a transcription-dependent manner (*30*), these data suggest the existence of a cardiac-specific splicing factory involving RBM20. We propose that this structure might bring together in the 3D nuclear space loci that are found far away in their primary sequence or even on different chromosomes, thereby enhancing the efficiency of functional alternative splicing.

This study provides a comprehensive integration of Hi-C, RNA-seq and ATAC-seq across a defined time course of human hPSC differentiation into cardiomyocytes to investigate the interplay of local and global chromatin structure on gene regulation. These assays demonstrate dynamic regulation during differentiation and cell fate transitions. *In vitro* CMs exhibit similar nuclear organization to their *in vivo* counterparts. Active regions are associated both in *cis* and *trans* and have high chromatin accessibility and transcription. In contrast, heterochromatic, silent regions in hESCs are relatively accessible compared to differentiated cells but compact during differentiation coincident with increased long range Hi-C signal and a loss of ATAC-peaks. This is consistent with recent electron micrograph studies on chromatin that show heterochromatic regions are more densely packed relative to euchromatic regions in differentiated cells (*31*). While the scope of the current study focused on large scale genome organization during normal differentiation, future studies will examine how this process is disrupted in congenital heart disease, and other diseased states.

## Acknowledgements

All data for Hi-C, RNA-seq and ATAC-seq is available on Gene Expression Omnibus accession number GSE106690. Thank you to members of the Murry lab for helpful discussions during the development of this project and especially to Katie Mitzelfelt for experimental support. We thank members of the Shendure and Noble labs and in particular Ruolan Qiu and Kate Cook for assistance with Hi-C technology and analysis. Flow cytometry was done with assistance from UW Cell Analysis Facility. We would like to acknowledge the Mike and Lynn Garvey Cell Imaging Lab at UW and its director Dale Hailey for assistance with sample imaging and analysis. We thank members of the Birth Defects Research Laboratory for assistance in obtaining human heart tissue. Funding: PF is funded through Experimental Pathology of Cardiovascular Disease training grant, NIH T32 HL007312. AB is funded by an EMBO Long-Term Fellowship (ALTF 488–2017). This work is part of the NIH 4D Nucleome consortium (NIH U54 DK107979, to CEM, WSN and JS), with additional support from P01 GM081619, R01 HL128362, and the Foundation Leducq Transatlantic Network of Excellence (to CEM). The Birth Defects Research Laboratory is supported by NIH grant R24 HD000836.

## Supplementary Materials

Materials and Methods

Table S1 - S6

Fig S1 – S7

References 32–48

